# Cryo-EM structure of the human TACAN channel in a closed state

**DOI:** 10.1101/2021.08.23.457436

**Authors:** Xiaozhe Chen, Yaojie Wang, Yang Li, Xuhang Lu, Jianan Chen, Ming Li, Tianlei Wen, Ning Liu, Shenghai Chang, Xing Zhang, Xue Yang, Yuequan Shen

## Abstract

TACAN is an ion channel involved in sensing mechanical pain. It has recently been shown to represent a novel and evolutionarily conserved class of mechanosensitive channels. Here, we present the cryoelectron microscopic structure of human TACAN (hTACAN). hTACAN forms a dimer in which each protomer consists of a transmembrane globular domain (TMD) that is formed of six helices and an intracellular domain (ICD) that is formed of two helices. Molecular dynamic simulations suggest that a putative ion conduction pathway is located inside each protomer. Single point mutation of the key residue Met207 significantly increased the surface tension activated currents. Moreover, cholesterols were identified at the flank of each subunit. Our data show the molecular assembly of hTACAN and suggest that the wild type hTACAN is in a closed state, providing a basis for further understanding the activation mechanism of the hTACAN channel.

## Introduction

Mechanotransduction, the conversion of mechanical stimuli into electrochemical signals, plays a vital role in all kinds of life forms. The primary sensors that react quickly to mechanical signals are mechanosensitive ion channels (MSCs) [1–3]. A variety of MSC families, including mechanosensitive channel large conductance (MscL) [4], two-pore potassium (K2P) channels [5], piezos [6], osmosensitive calcium-permeable cation (OSCA) channels [7] and transmembrane channel-like protein 1/2 (TMC1/2) [8], have been identified. Each has a unique structure and function [1]. These channels are involved in several physiological functions, such as hearing, touch, pain and proprioception [2, 3, 9].

Pain is a biological warning signal that is detected by nociceptors, which are activated by multiple environmental factors, and is sensed by the opening of several types of MSCs, including piezos in mammals [10, 11]. It was shown that piezos are associated with the sensation of touch and mechanical allodynia [6]. A recent study reported that the transmembrane protein 120A, referred to as TACAN, is a novel ion channel responsible for sensing acute mechanical pain [11] and is involved in mechanically sensitive nociception and mechanical pain perception currents in mice [12].

The TACAN is conserved in vertebrates and shares no sequence homology with other membrane proteins. It was originally identified as a nuclear envelope protein [13] and may be involved in adipocyte differentiation [14]. TACAN contains 343 amino acids and consists of six transmembrane helices with intracellular amino and carboxyl termini [11]. To better understand its domain organization and channel assembly, we determined the structure of the human TACAN (hTACAN) channel using single-particle cryo-electron microscopy (cryo-EM).

## Results

### Structure determination

Structural studies of the full-length hTACAN channel were performed by cryo-EM single particle analysis. The purified hTACAN channel protein showed a molecular weight of 80 kDa (approximately corresponding to a dimer) on an analytical size exclusion column (Fig S1A). The peak fraction was then collected and concentrated for cryo-EM sample preparation (Fig S1B). The 2D class average suggested that the protein is a dimer with 2-fold symmetry (Fig S1C). Subsequent 3D reconstructions of the hTACAN channel were determined at an overall resolution of 3.66 Å (Fig S1D–S1G). The electron density map for most amino acid side chains of the hTACAN channel was clearly resolvable (Fig S1H); thus, an atomic model with good stereochemistry was built. The final model includes most amino acids except the N-terminal region (aa 1-7), disordered loop (aa 249-263) and C-terminal region (aa 336-343).

### Overall structure

The overall structure of the hTACAN channel assembles as a dimer with a 2-fold rotation axis perpendicular to the membrane bilayer (Fig 1A–1C). It forms a two-layer architecture: the upper transmembrane part (termed the TMD) consistsC of two globular domains, and the lower cytosolic part (termed the ICD) consists of domain-swapped four-helix bundles (Fig 1D). Two sterol molecules were identified at two sides of the dimer (Fig 1D). Each globular domain is buried in the membrane bilayer and contains six transmembrane helices termed TM1-TM6 and a small anchoring helix termed A0 (Fig 1E, 1F and S2A). The anchoring helix is reminiscent of the anchor domain in piezo channels [15–17]. It is located in the dimeric interface and may contribute to the stabilization of the dimer. The ICD of each protomer contains two helices termed H1 and H2. H1 forms an extralong helix extending to the other protomer, resulting in the N-terminal half of H1 in one protomer interacting with the C-terminal half of H1 and the entire H2 of the other protomer.

**Fig. 1.**
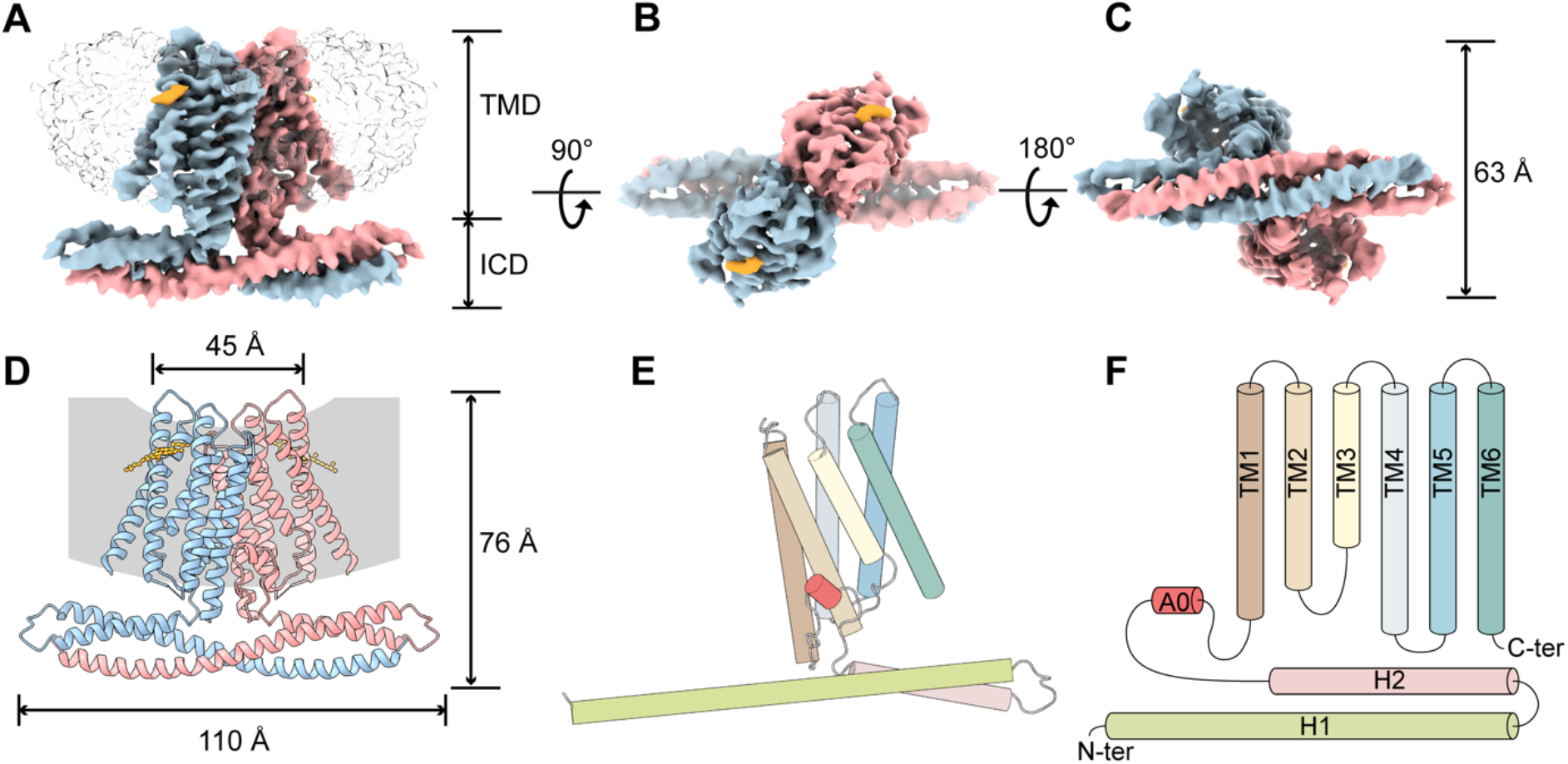
Structure of hTACAN. **A**-**C**, Side (**A**), top (**B**) and bottom (**C**) views of the cryo-EM density map. The two protomers are colored blue and pink. The sterol densities are colored yellow. Two globular domains of hTACAN are buried in detergent micelles represented by the transparent gray map. **D**, **E**, Cartoon representation of the hTACAN dimer (**D**) and the hTACAN protomer (**E**). The cholesterol model was drawn to represent the sterol for clarity. **F**, Schematic illustration outlining the protein secondary structures of one hTACAN protomer.

Compared with previously published mechanosensitive ion channels, the hTACAN channel shares a dimeric architecture with the *Arabidopsis thaliana* OSCA (AtOSCA) channel [18, 19] despite having no sequence homology to this protein. However, the topologies of the protomers of the hTACAN channel and AtOSCA channel are completely different (Fig S2B and S2C). Each protomer of the AtOSCA channel contains 11 TMs, while the hTACAN channel has only 6. In addition, the beam domain [15, 16] or the beam-like domain [18] which connects to pore-lining helices and plays an important role in gating the piezo channel or the AtOSCA channel, is absent in our hTACAN channel structure. Moreover, the intersubunit cleft of the AtOSCA channel is sufficiently wide to be filled with phospholipid molecules [18] while the intersubunit cleft of the hTACAN channel is too narrow to accommodate phospholipid molecules, as shown by molecular dynamics simulations (Fig S3). These results suggest that the hTACAN channel has a novel mechanogating mechanism.

### Intersubunit interactions

Interactions of the two hTACAN channel protomers occur at both the TMD interface and the ICD interface (Fig 2A). Due to the presence of the 2-fold symmetry axis, all interactions between the two protomers are reciprocal. The TMD interface can be further divided into the TM/TM interface (Fig 2B), loop A0-TM1/loop A0-TM1 interface (Fig 2C) and TM/A0 interface (Fig 2D). The TM/TM interface contains one hydrophobic pair interaction between two prolines (P211) of the TM3 from each protomer. The loop A0-TM1/loop A0-TM1 interface is involved in the hydrophobic interactions between residue V116 of one protomer and residue V118 of another protomer. The TM/A0 interface includes three hydrogen bonds mediated by the residue R178 of the TM1 from one protomer and three main chain oxygen atoms of S110, V112 and L113 of the other protomer. The ICD interface contains extensive interactions between two H1 helices from each protomer (Fig 2E) and between the helix H1 from one protomer and the helix H2 from another protomer (Fig 2F).

**Fig. 2.**
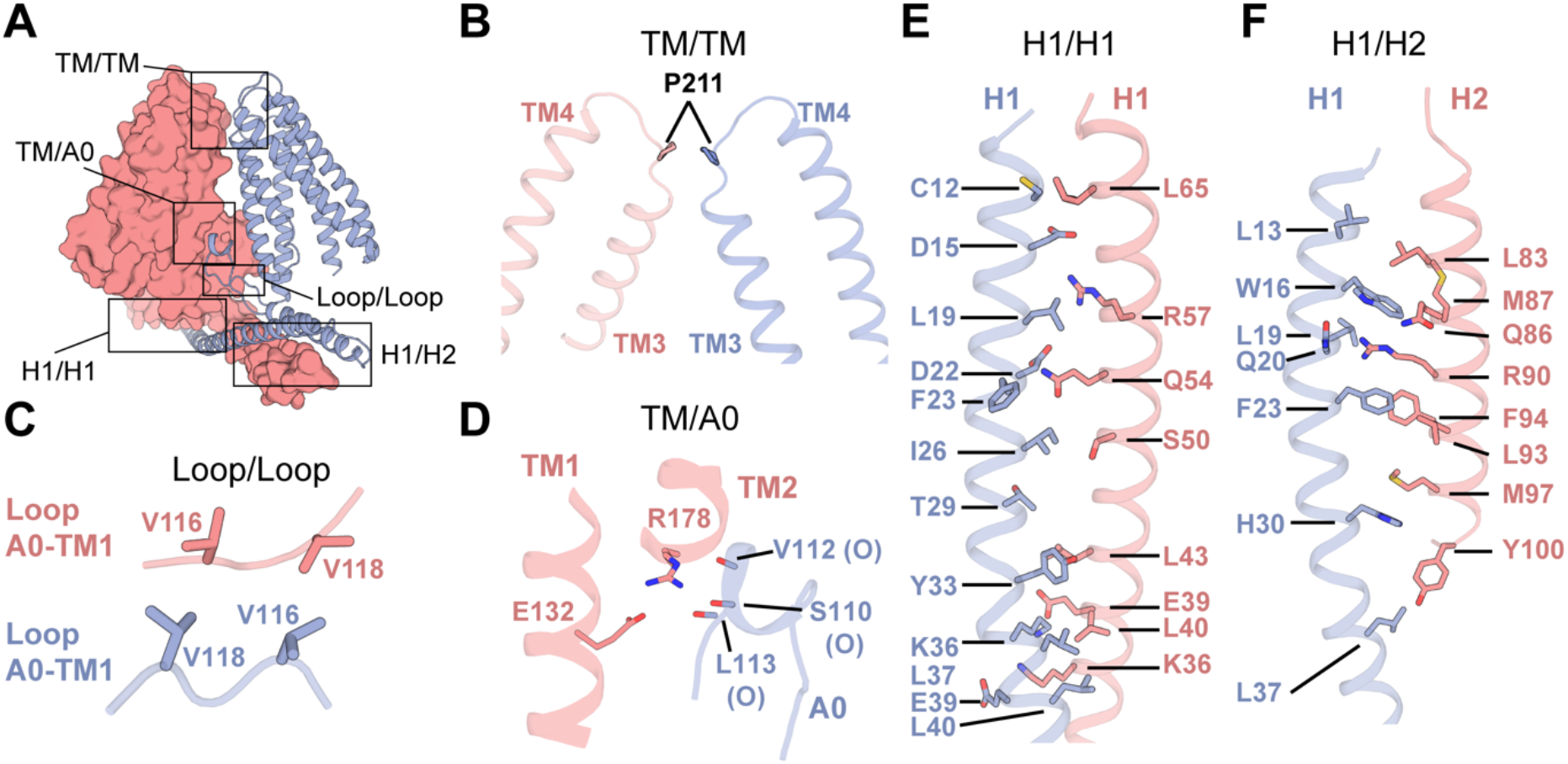
Intersubunit interfaces. **A**, Side view of the overall illustration of the intersubunit interactions. Two protomers are shown in the surface representation (pink) and cartoon representation (blue). **B**, The interaction between two TM3 helices at residue Pro211. **C**, Interactions between two loops. The loop A0-TM1 denotes that the loop is located between the A0 helix and the TM1 helix in each protomer. **D**, Interactions between the TM2 helix and the A0 helix. The side chain of R178 forms hydrogen bonds with main chain oxygen atoms from three residues S110, V112 and L113. **E**, **F**, Interactions between two H1 helices (**E**) and between helix H1 and helix H2 (**F**). It is noted that all these interactions described in (**C**), (**D**), (**E**), and (**F**) are reciprocal.

To further confirm the intersubunit interactions, we performed a coimmunoprecipitation (co-IP) experiment. We constructed three truncation mutants: delta-halfH1 (residues 43-343), delta-H1 (75-343) and delta-ICD (101-343). C-terminal FLAG or GFP tags were added to these mutant proteins. Then, the mutant plasmids with two different tags were cotransfected into HEK293T cells for co-IP. Our results showed that the deletion of H1 significantly attenuated the dimeric assembly, suggesting that the dimeric hTACAN channel requires the integrity of the ICD (Fig S4).

### Ion-conduction pathway

The surface map of the hTACAN dimer shows that there is a large fenestration in the middle of the intersubunit cleft from the side view (Fig 3A). Protein atoms occupied spaces around the fenestration. Therefore, the intersubunit cleft is unlikely to act as an ion conducting pathway. Instead, a vestibule toward the intracellular side was identified. This vestibule is positively charged in the protomer according to the surface potential map (Fig 3B and 3C). However, at the extracellular side, it is difficult to find an opening to connect with this large vestibule to act as a putative ion conducting pathway in our current structure. We therefore conducted molecular dynamics (MD) simulations. The 200-ns simulation did not produce a continuous distribution of water passages within each subunit (Fig 3D), indicating that our hTACAN channel structure represents a nonconducting state. To investigate the conformational changes occurring in the hTACAN channel in response to mechanical stress, different surface tensions were applied to the lipid bilayer during the atomic MD simulations. After a surface tension of 35 mN/m was applied to the system, water molecules were able to spontaneously form a continuous channel from the extracellular side to the intracellular side in each protomer (Fig 4A). The corresponding ion conduction pathways were calculated by the HOLE program [20] (Fig 4B). The restriction region of the ion conduction pathway was composed of three residues: N165 of TM2, M207 of TM3 and F223 of TM4 (Fig 4C). Specifically, the side chains of M207 and F223 undergo large conformational changes before and after surface tension in MD simulations. As shown in Fig 4D, in response to the surface tension, this restriction region increases from the narrowest radius of 0.27 Å to approximately 1.24 Å, sufficient to allow the passage of some dehydrated cations such as Ca^2+^, Mg^2+^, Na^+^, etc. (Fig 4D). To verify the ion conduction pathway, we conducted electrophysiological studies. GFP only (mock), GFP-tagged wild-type hTACAN and GFP-tagged mutant M207A plasmids were transiently transfected into COS7 cells, cell-attached configuration mode was then applied. A cell-surface biotinylation assay showed that the surface expression levels of the mutant M207A were similar to those of the wild type (Fig S5A). Comparing with mock, the currents measured in response to different surface tensions for wild type hTACAN was no marked difference, while the currents for the mutant M207A showed about 6-fold increasement (Fig 4E and S5B). Therefore, it seems that the wild type hTACAN channel is in a closed state and other unknow factors may be required to activate the hTACAN channel.

**Fig. 3.**
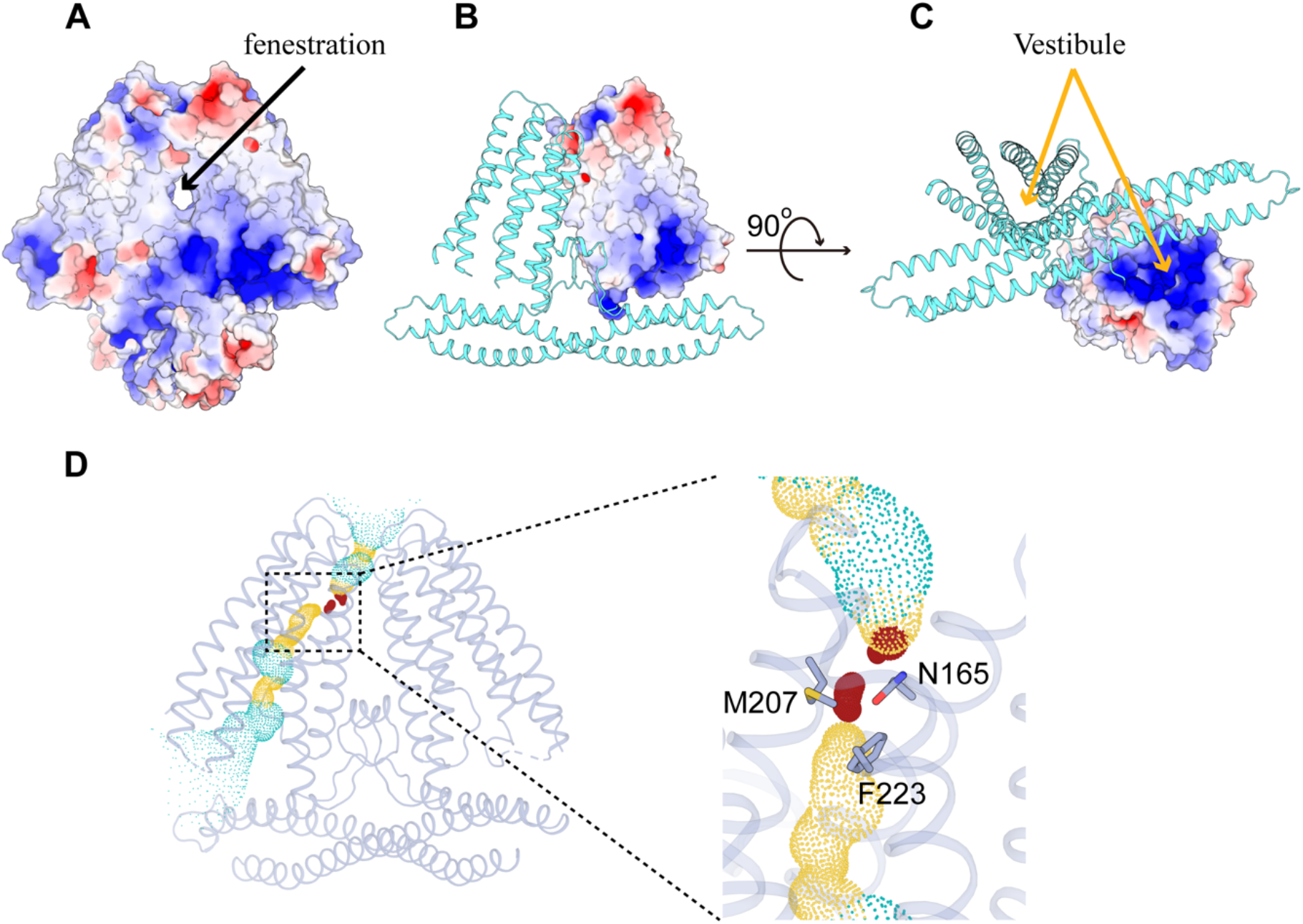
Ion conduction pore of hTACAN. **A**, Surface potential map of the hTACAN channels. The hTACAN channels in the surface representation are colored according to the electrostatic surface potentials from −5 to 5 kT/e (red to blue). **B**, **C**, Vestibule inside one protomer of the hTACAN channel. The TM domain of one protomer is shown in the surface presentation. **D**, Potential ion conduction pore inside one protomer. The constriction region of the pore is magnified on the right side. The dots colored blue, red and orange represent the pathways of the pore. Among colors, the red dots denote that the pathway is blocked in this area.

**Fig. 4.**
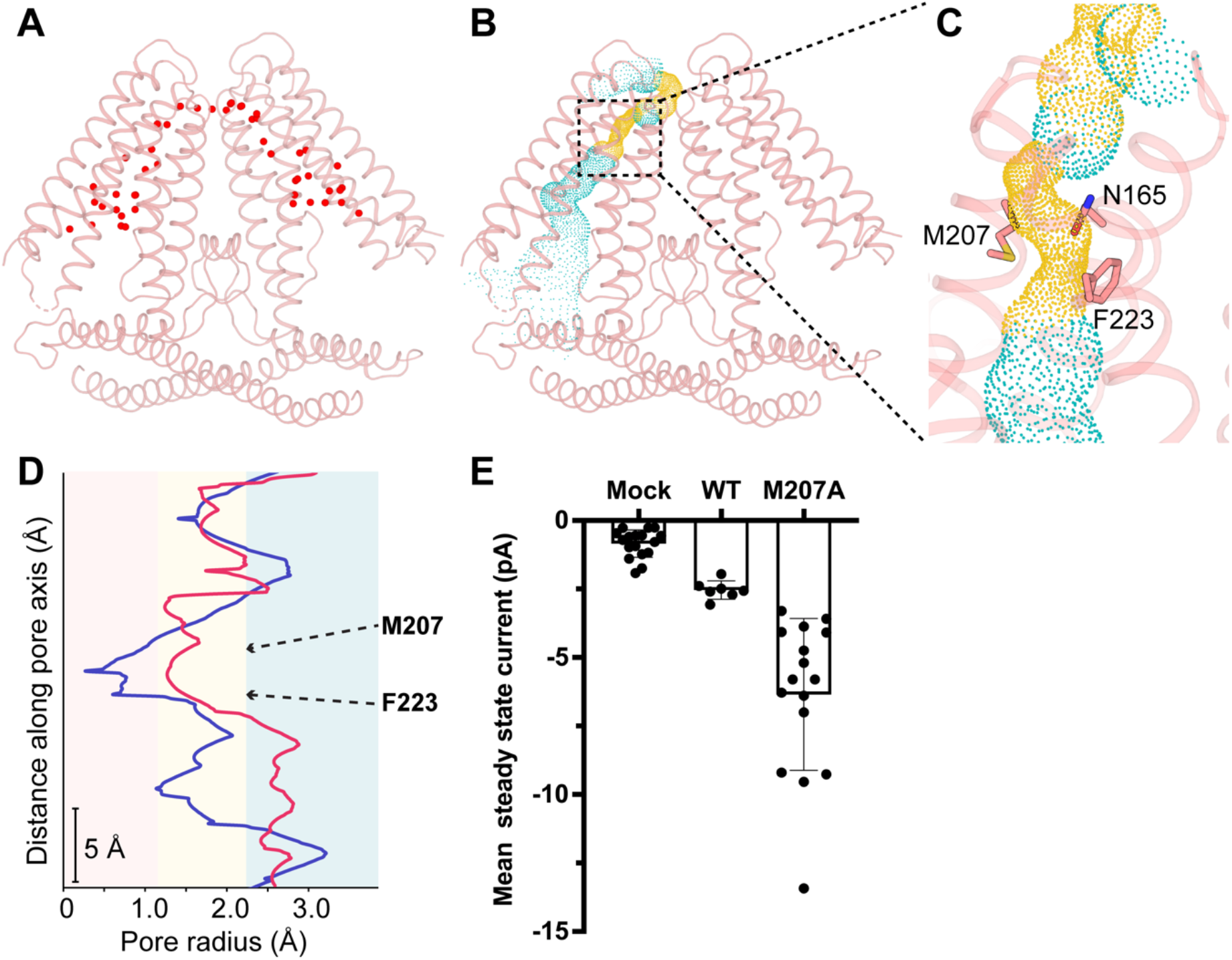
Ion conduction pore. **A**, Continuous water channel within the hTACAN dimer in response to a surface tension of 35 mN/m by MD simulations. The red dots denote the water molecules. The cartoon representation of the hTACAN dimer is colored pink. **B**, Side view of the equilibrated structure after 200-ns MD simulations with 35 mN/m surface tension. The putative pore is indicated as orange and blue dots. **C**, Zoom view of the restricted area in the pore. Side chains of two residues, M207 and F223, formed the hydrophobic gate. **D**, The pore radius profiles of TACAN before (blue) and after (red) 200-ns MD simulations with surface tension. **E**, Mean currents for individual cells expressing wild-type hTACAN (n=7) or the mutant M207A (n=16) or the mock (GFP only, n=17). The error bars denote the SEM.

**Fig. 5.**
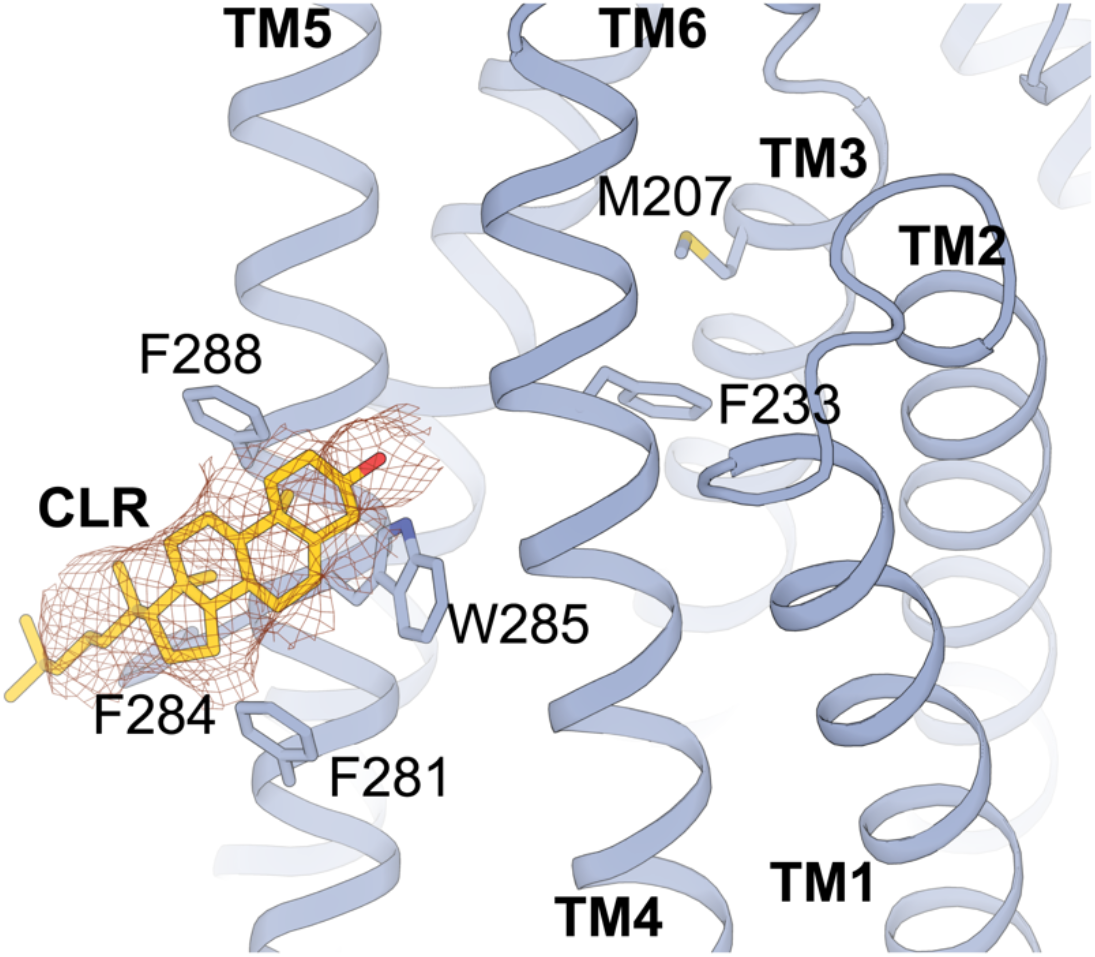
Cholesterol binds to the hTACAN. The binding site of cholesterol (CLR) on TM5. TACAN is shown in blue and the CLR molecule is shown in yellow. The electron density map of the CLR molecule is represented by a red meshed line, and the side chains of residues involved in CLR binding (F281, F284, W285 and F288) are shown as stick models. The residues involved in pore formation (M207 and F223) are also shown in a stick model to show their positions relative to the CLR molecule.

### Cholesterol binding

In the hTACAN channel structure, we observed a robust density that fits well with the cholesterol molecule at the flanks of the two protomers (Fig 4). Mass spectrometry indeed identified the major peak of the molecule weight of 387 Dalton, which is corresponding with the cholesterol (Fig S6A), from the purified hTACAN protein sample. No other lipid molecules were identified in our experiment. The hydrophobic steroid nucleus of the cholesterol is stabilized by multiple residues with bulky hydrophobic side chains, including F281, F284, W285 and F288 of TM5. We then made the mutant hTACAN channel F284A-W285A-F288A. The subsequent mass spectrometry analysis of the purified mutant protein sample did not find the cholesterol peak (Fig S6B). Although cholesteryl hemisuccinate (CHS, cholesterol mimicking detergent) was used in protein sample purification, we did not find the peak corresponding with the CHS molecule in the mass spectrometry analysis. Moreover, Phospholipid molecules have been shown to be bound in multiple mechanosensitive channels and to be involved in their activation [1]. However, phospholipid molecules have not been found in our current TACAN structure. These results suggest that cholesterol is a specific binding molecule for the hTACAN channel. The functional role the cholesterol played in hTACAN channel needs to be further clarified.

## Discussion

Our studies described here showed that hTACAN represents a structurally novel class of ion channels. Although its topology is different, the dimeric architecture and ion conduction pathway of hTACAN are reminiscent of those of AtOSCA channels, suggesting that these two tension-activated mechanosensitive channels may share some commonalities upon activation. Importantly, cholesterols were identified in the TACAN channel, which, to the best of our knowledge, has never been found in any known mechanosensitive channel.

Recently published literatures argued that the TACAN may be a coenzyme A-binding protein rather than a mechanosensitive channel [21–23]. They showed that a coenzyme A molecule is bound in the vestibule of the membrane domain of hTACAN and also the expression of hTACAN was not sufficient in mediating mechanosensitive currents in cells. However, these results may suggest that the wild type hTACAN channel is in a closed state due to the binding of coenzyme A to block the ion conduction. Additional subunits may be required to release the coenzyme A in order to activate the hTACAN channel. In other words, there is a possibility that functional hTACAN channel contains multiple subunits. Mechanosensitive channels can be made up of several subunits. For example, in hair cell, mechanoelectrical transduction channels consist of transmembrane channel-like 1 and 2 (TMC1/2), transmembrane inner ear (TMIE), transmembrane proteins lipoma HMGIC fusion partner-like 5 (TMHS/LHFPL5), and may be more [24].

Our results seem to be consistent with this hypothesis. In our hTACAN structure, we did not find any densities corresponding with the coenzyme A, indicating that the binding of coenzyme A to hTACAN may be dynamic. Moreover, our MD and electrophysiology data showed that the residue M207 seems to act as a gate to block the ion conduction in wild type hTACAN channel. The single point mutation of M207A indeed made the channel more permeable.

Taken all together, we propose that other unknow factors and cholesterols may be involved in the movement of TM5 and TM4 helices, thus eliciting the shift of side chains of two hydrophobic residues (F223 and M207) to open the channel (Fig S7). Further studies are required to understand the detailed conformational transition during activation, but our structure of hTACAN in a closed state will be helpful for the future investigation of the hTACAN channel in a constitutively active state and contribute to the development of a novel class of painkillers.

## Materials and Methods

### Plasmid

The gene encoding *Homo sapiens* TACAN (Uniprot: Q9BXJ8) was synthesized (Genewiz, Inc.) and cloned into pEG-BacMam vector, with a 3C protease site and GFP tag at c-terminal. This plasmid was used to generate Baculovirus using the Bac-to-Mam Baculovirus expression system (Invitrogen).

### Protein expression and purification

The plasmid expressing wild-type hTACAN was transfected into DH10Bac cells to produce Bacmid. Adherent sf9 cells grown at 27 ° C were transfected with purified Bacmid to produce P1 virus. The P1 virus then infected the sf9 suspension cells at a 1:80 (v/v) ratio to produce P2 virus. The HEK293S-GnTi^−^ cells were cultured in Freestyle 293 medium (Thermo Fisher Scientific) supplemented with 5% CO_2_ at 37 ° C. The cells were transfected with P2 virus of hTACAN at 1:50 (v/v) ratio when the cell density reached 2×10^6^ cells per ml. After 12h, sodium butyrate was added to the cell culture at a final concentration of 10 mM and the temperature was adjusted to 30 ° C to facilitate protein expression. After 60h, suspension cells were collected and store at −80 ° C.

For protein purification, 1.5 liters of cells were resuspended in buffer containing 20 mM Tris-HCl pH 8.0, and 200 mM NaCl. The suspension was supplemented with 0.5% (w/v) lauryl maltose neopentyl glycol (LMNG, Anatrace), 0.1% (w/v) cholesteryl hemisuccinate tris salt (CHS, Anatrace) and 1× protease inhibitor cocktail (Tagetmol). After lysed at 4 ° C for 2h, the sample was ultracentrifuged for 50 min at 130,000×g and the supernatant was incubated with sepharose resin conjugated with anti-GFP nanobody at 4 ° C by gentle rotation for 2.5h. The resin was washed with buffer containing 20 mM Tris-HCl pH 8.0, 200 mM NaCl, 0.01% (w/v) LMNG with 0.001% (w/v) CHS and 0.0033% (w/v) glycol-diosgenin (GDN, Anatrace) with 0.00033% (w/v) CHS. Afterwards, the hTACAN protein was eluted by incubating resin with rhinovirus 3C protease at a ratio of 1:40 (v/v) to cleave GFP tag overnight. The eluent was then concentrated to 1 mL using a 100 kDa cut-off concentrator (Millipore) and further purified with superose 6 10/300 GL (GE Healthcare), which was equilibrated with buffer containing 20 mM Tris-HCl pH 8.0, 200 mM NaCl, 0.00075% (w/v) LMNG and 0.00025% (w/v) GDN. The peak fractions were pooled and concentrated to 25 mg/mL for Cryo-EM grid preparation.

### Cryo-EM sample preparation

An aliquot of 2 μL purified hTACAN was placed on glow-discharged holey carbon film gold grid (Protochips, CF-2/1, 200 mesh). The grid was blotted at 8 ° C and 100% humidity and plunge-frozen in liquid ethane using FEI Vitrobot Mark IV (Thermo Fisher Scientific). Cryo-EM data was collected on a Titan Krios electron microscope operated at an acceleration voltage of 300 kV with the defocus range of −1.0 μm to −2.0 μm using SerialEM software [25]. Micrographs were recorded by a Gatan K2 Summit direct electron detector in super-resolution mode with a pixel size of 0.507 Å (a physical pixel size of 1.014 Å) with a dose rate of 7.2 electrons per pixel per second for a total exposure time of 8 s, resulting in an image stack with 40 frames.

### Image Processing

A total of 2,585 image stacks were collected and subjected to beam-induced motion correction using MotionCor2 [26] with binning to physical pixel size. The contrast transfer function (CTF) parameters for each micrograph were estimated using CTFFIND4 [27] on the basis of summed images without dose-weighting. Micrographs of low-quality were excluded manually according to the CTF power spectra and summed images in Relion-3.1 [28]. A total of 682,480 particles from 2,068 micrographs were picked semi-automatically using Gautomatch (http://www.mrc-lmb.cam.ac.uk/kzhang/). Two rounds of 2D classification were performed to exclude noise and poorly defined particles using Relion-3.1. The well-defined subsets accounting for 369,479 particles were extracted in Relion-3.1 and then used to generate an initial model through the *Ab initio* reconstruction in CryoSPARC-2 [29]. The initial model was low passed to 30 Å resolution for the first round of 3D classification in Relion-3.1. After 2 rounds of 3D classification (3 expected classes in each round), 64,446 particles were selected and subsequently subjected to 3D refinement and Bayesian polishing in Relion-3.1. The polished particles were then imported to CryoSPARC-2 for heterogenous refinement and non-uniform refinement, a final map at 3.66 Å resolution was produced from 58,843 particles with C2 symmetry imposed. The final resolution was estimated based on gold-standard Fourier shell correlation (0.143 criteria) after correction for the use of masks. The local resolution map was calculated using ResMap [30] and exhibited using UCSF Chimera [31]. Data collection and reconstruction workflow are shown in fig. S1.

### Model building and refinement

The atomic model of hTACAN was manually built in COOT software [32], sequence assignment was mainly guided by secondary structure prediction results from PSIPRED Workbench [33] and visible densities of residues with bulky side chains (Trp, Phe, Tyr and Arg) and refined using the real-space refinement in Phenix [34]. The quality of the final model was validated using the module ‘Comprehensive validation (cryo-EM)’ in Phenix. Refinement and validation statistics are provided in Extended Data Tab. 1. The figures showing structures were generated by the UCSF ChimeraX package [31], UCSF Chimera and PyMol (Schrödinger, USA).

### Co-Immunoprecipitation

Transfected HEK293T cells were washed twice with Tris-buffered saline (TBS) and lysed in buffer containing 20 mM Tris-HCl (pH 8.0), 200 mM NaCl, 1% (w/v) DDM, 0.2% (w/v) CHS, and 1 × protease inhibitor cocktail (Roche). After incubation at 4 ° C for 1h, cell lysates were centrifuged at 120,000 ×g for 40 min at 4 °C. The supernatant was divided into two equal parts. One part was incubated with GFP-Trap (20 μL slurry, ChromoTek), and the other was incubated with anti-flag affinity beads (20 μL slurry, BioLegend) for 2-3h at 4 ° C. The beads were washed five times with buffer containing 20 mM Tris-HCl (pH 8.0), 200 mM NaCl, 0.025% (w/v) DDM, 0.005% (w/v) CHS and eluted with 40 μL gel-loading buffer. Then, 5 μL of the eluate was loaded onto 12% SDS-PAGE gels and transferred to a PVDF membrane for western blot analysis.

### Cell surface biotinylation assay

The transfected HEK293T cells were washed twice with ice-cold 1×PBS and then treated with 1 mg/mL biotinamidohexanoic acid 3-sulfo-N-hydroxysuccinimide ester sodium salt in PBS for 40 min at 4 °C. Then, the cells were washed with an ice-cold buffer containing 20 mM Tris-HCl (pH 8.0) and 200 mM NaCl and incubated at room temperature for 15 min. Cells were collected by scraping and lysed in buffer containing 20 mM Tris-HCl (pH 8.0), 200 mM NaCl, 1% (w/v) DDM, 0.2% (w/v) CHS, and 1 × protease inhibitor cocktail. After incubation at 4 ° C for 1h, cell lysates were centrifuged at 120,000 ×g for 40 min at 4 °C. Each sample of supernatant was incubated with 40 μL of 50% NeutrAvidin agarose at 4 °C overnight. The beads were then washed three times with lysis buffer, eluted by incubation with SDS-PAGE sample buffer at 37 °C for 30 min and analyzed by SDS-PAGE followed by western blotting.

### Electrophysiology

COS7 cells were transiently transfected with the wild type or mutants TACAN-GFP for 24 to 48 h at 37 °C with 5% CO_2_. Transfected cells were then digested by trypsin to be plated onto 35-mm dishes for cultivation for at least 3 h before electrophysiology. For the cell-attached recordings, the membrane voltage was held at −80 mV. The electrode holder was directly connected to a high-speed pressure clamp and pressure-vacuum pump (HSPC, ALA Scientific). The pressure pulse was delivered through the recording electrode directly. Each cell membrane patch was subjected to 16 sweeps (500 ms for each sweep) of incrementally increasing negative pressure from 0 mmHg to −150 mmHg at −10 mmHg increments. Cell-attached currents were amplified with an Axopatch 200B and digitized with a Digidata 1550A system (Molecular Devices, Sunnyvale). All currents were sampled at 10 kHz and low-pass filtered at 0.5 kHz through the pCLAMP software. For cell-attached recordings, both the extracellular bath solution and the pipette solution consisted of the following (in mM): NaCl 140, KCl 5, CaCl_2_ 2, MgCl_2_ 2, HEPES 10 and glucose 10 (pH adjusted to 7.4 with NaOH, 305 mOsm).

### Coarse-grained MD simulations

Coarse-grained MD simulations were performed by using the Gromacs v.2020.2 program [35]. The protein coordinates of dimeric hTACAN were embedded in a 125 Å× 125 Å bicomponent bilayer composed of POPC and cholesteric (4:1) using the insane script [36]. The ElNeDyn elastic network [37] was applied to the protein. The system was solvated by the nonpolarizable water and neutralized at 0.15 M NaCl and 0.02 M CaCl_2_ concentrations. The size of the simulation box was 125 Å× 125 Å × 180 Å. The Martini force field version 2.2 was used for protein; and the version 2.0 force field was used for POPCs, cholesterols, and ions [38]. By using a velocity-rescaling thermostat [39] with coupling constants of τ_T_ = 1.0 ps, the temperature was maintained at 310 K. The pressure was controlled at 1 bar by the Berendsen thermostat [40] with a coupling constant of τ_P_=1.0 ps. The type of constrain applied to bonds was the LINCS algorithm [41]. A Verlet cutoff scheme was used, and the PME method [42] was applied to calculate long-range electrostatic interactions. Periodic boundary conditions were used in the x-, y- and z-directions. The time step was set to 20 fs. Three independent production phases were performed, and each production phase replicate was run for 1 μs.

### All-atom MD simulations

The final frames of the coarse-grained simulations were selected as the template to build the all-atom model system. The following steps were performed to provide the starting coordinates for all-atom MD: (i) the hTACAN-POPC-water system was transformed from the coarse-grained simulations above to the all-atom model using the backward.py protocol [43], (ii) the backward-generated hTACAN was replaced with the cryo-EM hTACAN structure after aligning the main chain of them, and (iii) the overlapped lipids and water molecules were removed through the operations in VMD [44]. In the hTACAN-POPC-water system, the protonation state for titratable residues of protein was determined using the H++ program [45], and the missing atoms were added automatically using the Tleap module in AMBER 20 [46].

All-atom MD simulations were conduct using AMBER 20 with the PMEMD engine [46]. The AMBER FF14SB force field [47] was used for proteins, and the AMBER lipid force field LIPID14 [48] was used for POPCs and cholesterols. A 12 Å cutoff was set for the nonbonded interaction. The SHAKE algorithm [49] integration was used to constrain the covalent bonds involving hydrogen atoms, and the Particle Mesh Ewald (PME) algorithm [42] was applied to treat long-range electrostatic interactions. First, the system was minimized for 10,000 steps. Second, each system was heated from 0 K to 310 K in 500 ps using the Langevin thermostat [50], and the proteins and lipids were constrained with a force constant of 50 kcal·mol^−1^·Å^−2^. Third, a series of equilibrations were performed for each system, and the lipids were equilibrated for 30 ns with the proteins constrained, followed by the 10-ns equilibration without any constraint applied for the entire system. Finally, a 200-ns production phase was performed for the all-atom system. The time step was set to 2 fs. The frames were saved every 5,000 steps for analysis.

Constant surface tensions was applied on the membrane surface (xy plane) by using the surface tension regulation in AMBER 20 [46], and the periodic boundary conditions and semi-isotropic pressure scaling were applied in the system. After the minimization, heat and equilibration phases mentioned above, three independent 200-ns production phases with surface tension values of 25, 30, and 35 mN/m, respectively, were performed. All other simulation parameters were set to the same as described above.

### Liquid chromatography mass spectrometry (LC-MS)

Purified hTACAN proteins were mixed with methanol and chloroform. The mixture was vortexed and incubated at room temperature for 15 min, then centrifuged at 14,000 ×g for 10 min. The aqueous layer on the top and the protein pellet layer in the middle were removed and discarded, the bottom chloroform layer was transferred into a new 1.5 mL tube and dried under vacuum. The dried extract was dissolved in 80 μL methanol with 0.1% formic acid for LC-MS analysis.

## Acknowledgements

We are grateful to thank Prof. B. Xiao at Tsinghua University for the help in electrophysiology studies, the technical assistance in the Center of Cryo-Electron Microscopy (CCEM), Zhejiang University on Cryo-EM for data acquisition, Dr. J. Lu, Dr. A. Li for the support of electron microscopy at Nankai University, and Dr. Q. Wang for the support of mass spectrometry facilities at Nankai University. This work was supported by National Key Research and Development Program of China (grant 2017YFA0504801 to Y.S.; 2017YFA0504803 and 2018YFA0507700 to X.Z.), National Natural Science Foundation of China (grants 91954119 and 31870736 to X.Y.; grants 32071231 and 31870834 to Y.S.) and the Fundamental Research Funds for the Central Universities (2018XZZX001-13) to X.Z.

## Author Contributions

X.C. did protein expression, purification, sample preparation and cryo-EM data collection; Y.W. performed electrophysiology experiment; Y.L. performed MD simulations; J.C., M.L. and W.T. performed protein expression and purification; X.L., S.C., X.Z. and X.Y. did cryo-EM data collection; X.Y. conducted cryo-EM reconstruction; N.L. performed mass spectrometry analysis. X.C., Y.W., Y.L., X.Y. and Y.S. analyzed the data, designed the study and wrote the paper. All authors discussed the results and commented on the manuscript.

## Data availability

Atomic coordinates have been deposited in the Protein Data Bank under accession number 7F6V. The cryo-EM density maps have been deposited in the Electron Microscopy Data Bank under accession number EMD-31482. Additional data related to this paper may be requested from the authors.

---

**Fig. S1.**
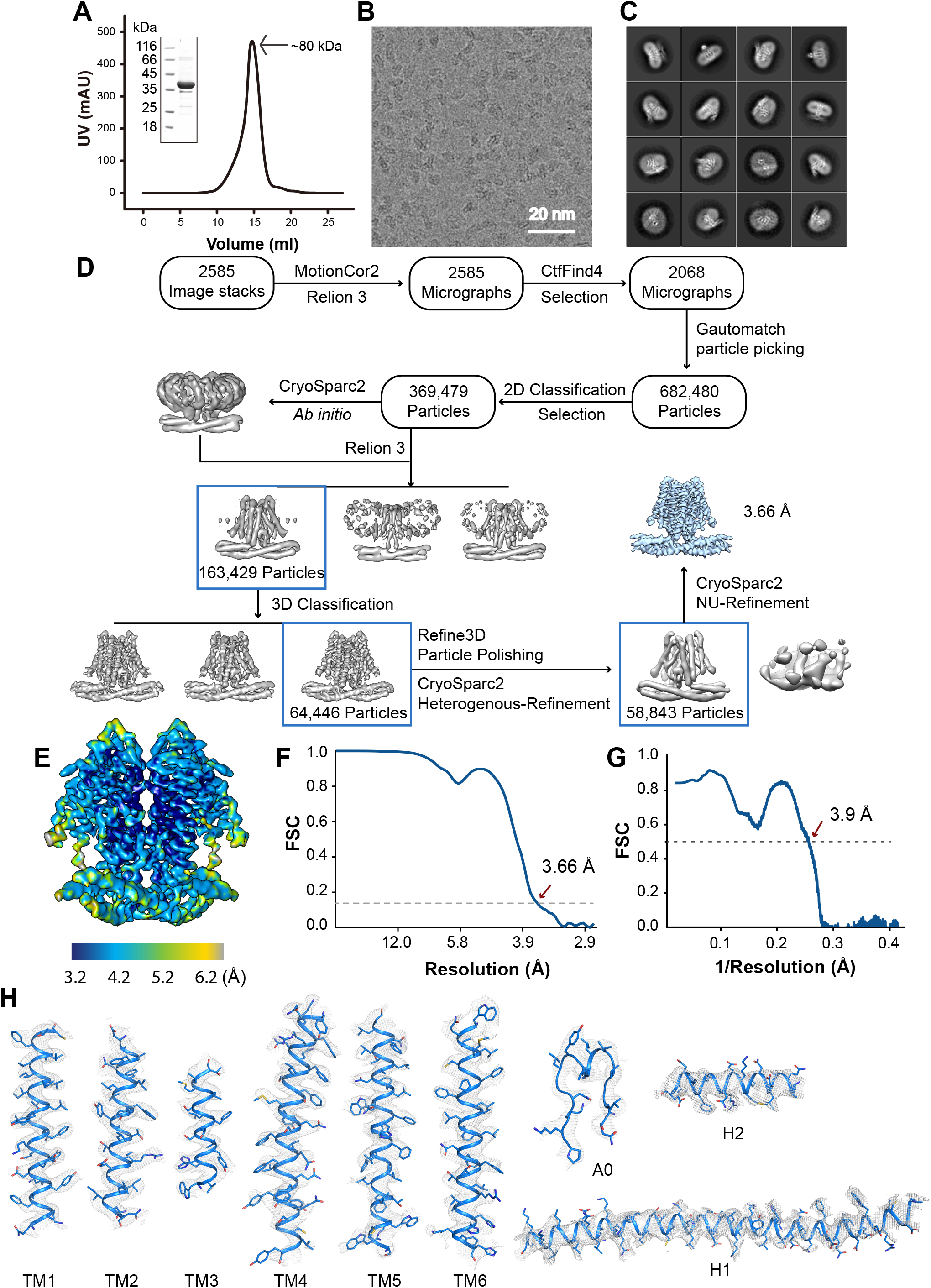
Structural determination of the hTACAN structure. **A**, Elution profile of hTACAN protein on the size-exclusion column Superose 6 10/300 GL. **B**, A drift-corrected cryoEM micrograph of hTACAN. **C**, Representative 2D class averages of hTACAN. **D**, Workflow of image processing, 3D reconstruction and structure refinement. **E**, Local resolution estimation from ResMap. **F**, Gold standard FSC curve of hTACAN estimated by cryo-SPARC v2. **G**, FSC curves for cross-validation between maps and models: model versus summed map. **H**, Representative densities of cryo-EM maps of hTACAN. Each secondary structural elements of hTACAN is shown in a blue cartoon. The densities are shown in gray.

**Fig. S2.**
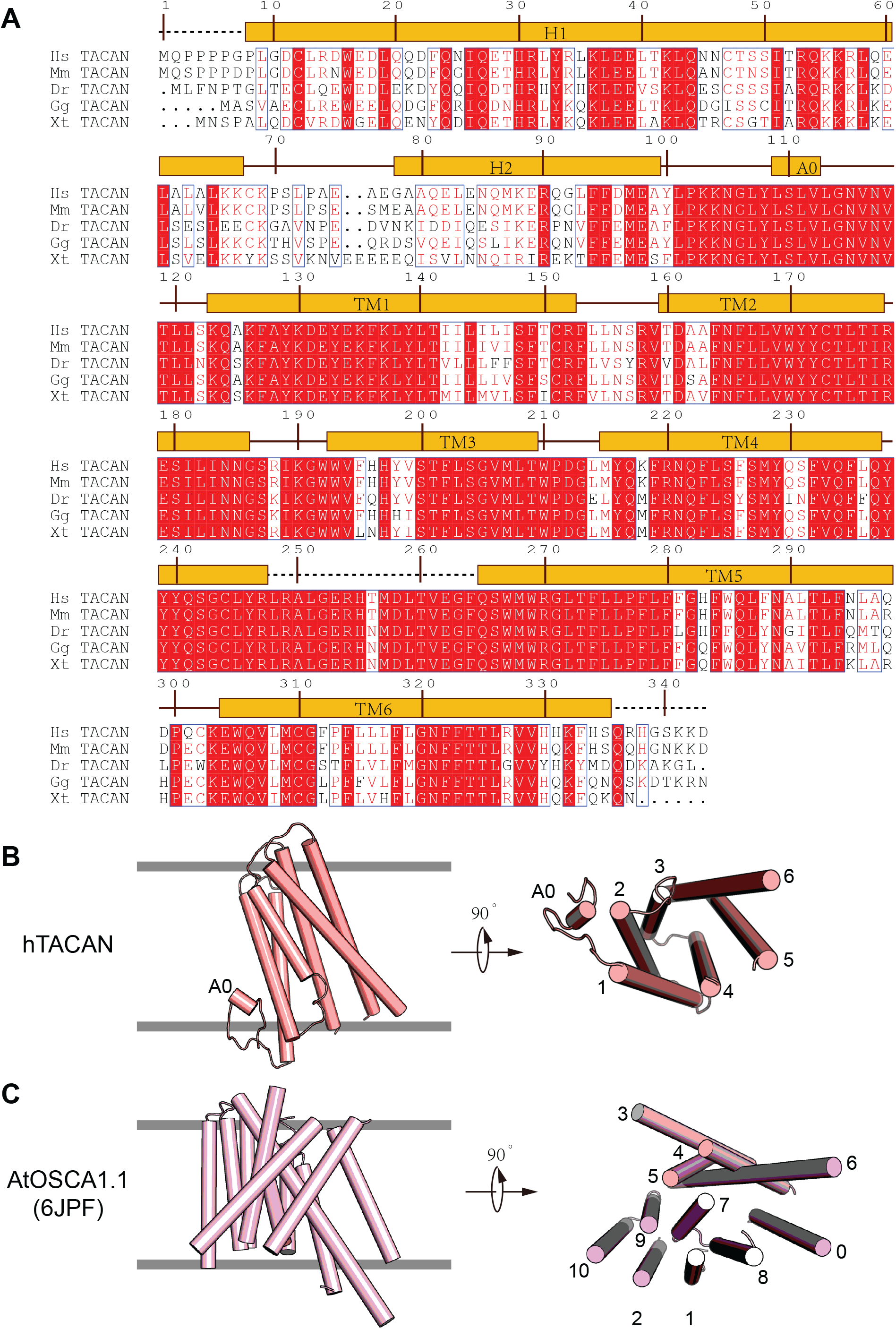
Structural comparison of hTACAN and AtOSCA1.1 channels. **A,** Sequence alignment of TACAN from various species. Sequence alignment of *Homo sapiens* TACAN (Hs TACAN, UniProt code Q9BXJ8), *Mus musculus* (Mm TACAN, UniProt code Q8C1E7), *Danio rerio* (Dr TACAN, UniProt code A3KNK1), *Gallus gallus* (Gg TACAN, UniProt code A0A1L1RMH9 and *Xenopus tropicalis* (Xt TACAN, UniProt code Q5FWV6). **B**, the transmembrane domain of the hTACAN channel. TM A0 was indicated, and other TM helices were labeled as numbers. **C**, Transmembrane domain of the AtOSCA1.1 channel (RCSB code 6JPF). The numbers (0-10) indicate each TM helix.B,

**Fig. S3.**
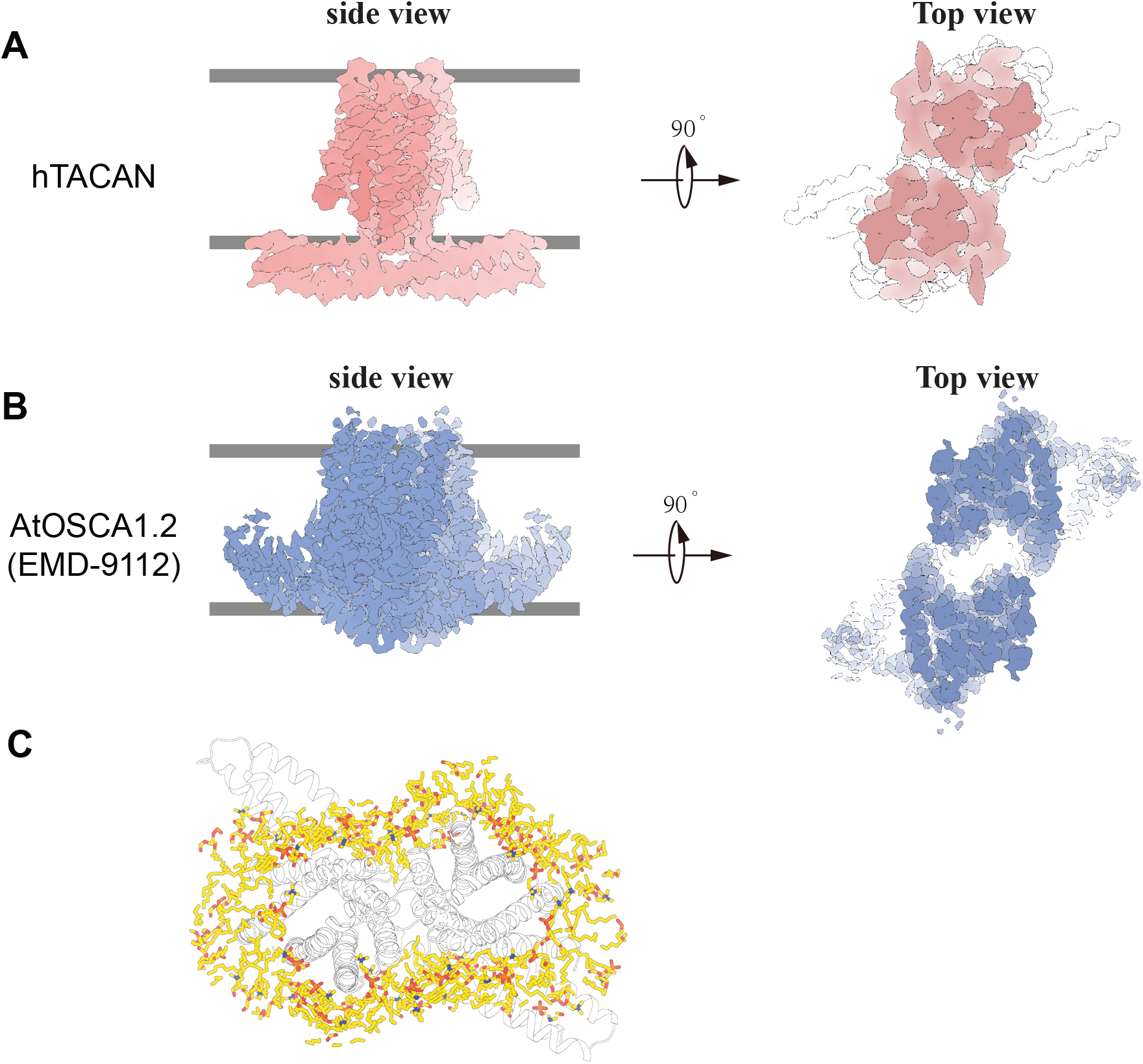
Intersubunit cleft of hTACAN and AtOSCA1.1 channels. **A**, Cryo-EM densities of the hTACAN channel in the side view and top view. The intersubunit cleft of the hTACAN channel is quite narrow. **B**, Cryo-EM densities of the AtOSCA1.2 channel (RCSB code EMD-9112) at two different views. The intersubunit cleft of the AtOSCA1.2 channel is much wider. **C**, Molecular dynamics simulations revealed the intersubunit cleft of the hTACAN channel to be surrounded but not be occupied by phosphatidylcholine molecules colored by yellow.

**Fig. S4.**
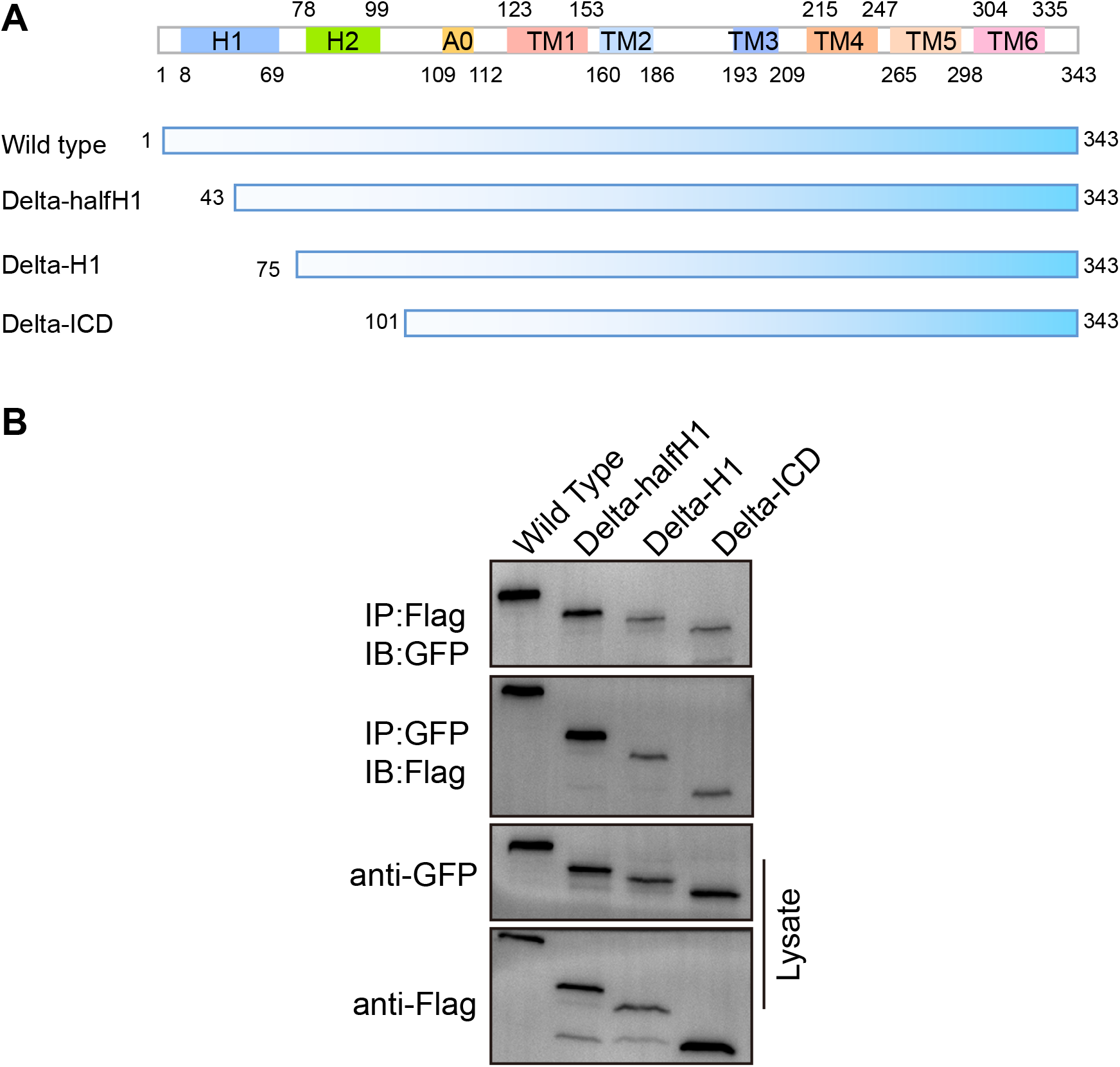
Dimer formation between various hTACAN mutants. **A**, Schematic drawing of hTACAN channel fragments. The secondary structure of the hTACAN channel is shown on the top. **B**, Co-immuno-precipitation experiment of various hTACAN mutants with a C-terminal FLAG tag or C-terminal GFP tag.

**Fig. S5.**
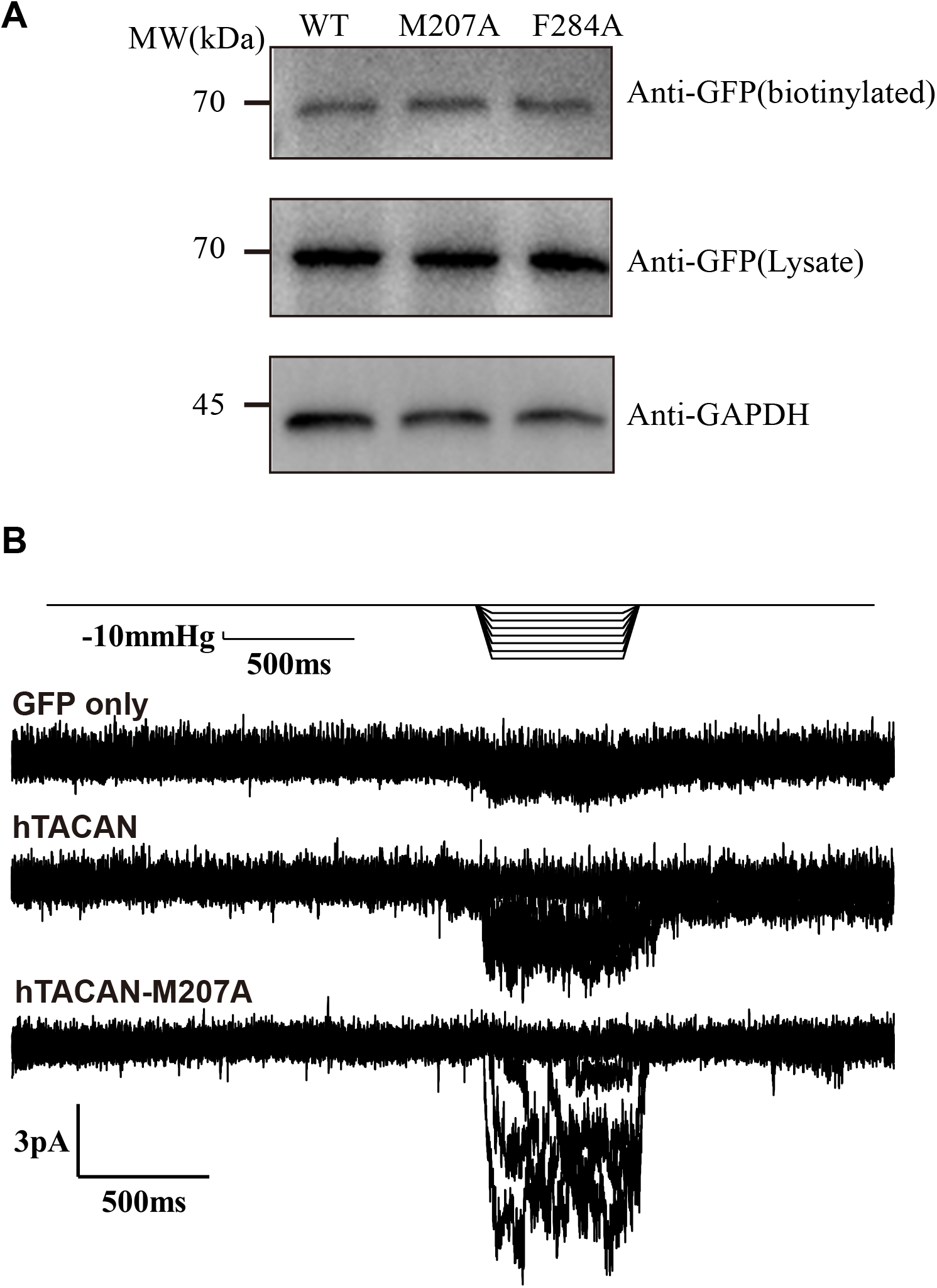
Currents of the COS7 cells in response to different surface tensions. **A**, Cell-surface biotinylation assay for wild type and mutant TACAN. Wild type and mutant TACAN were transiently transfected into COS7 cells, and cell-surface biotinylation assays were performed. The total expression level (lysate) and the surface amount (biotinylated) for wild type and mutant TACAN were similar. Glyceraldehyde 3-phosphate dehydrogenase (GAPDH) was used as an internal control. The experiment was repeated three times with similar results. **B**, Representative traces of one COS7 cell expressing the wild-type hTACAN channel or GFP only (mock) or hTACAN-M207A using the cell-attached configuration.

**Fig. S6.**
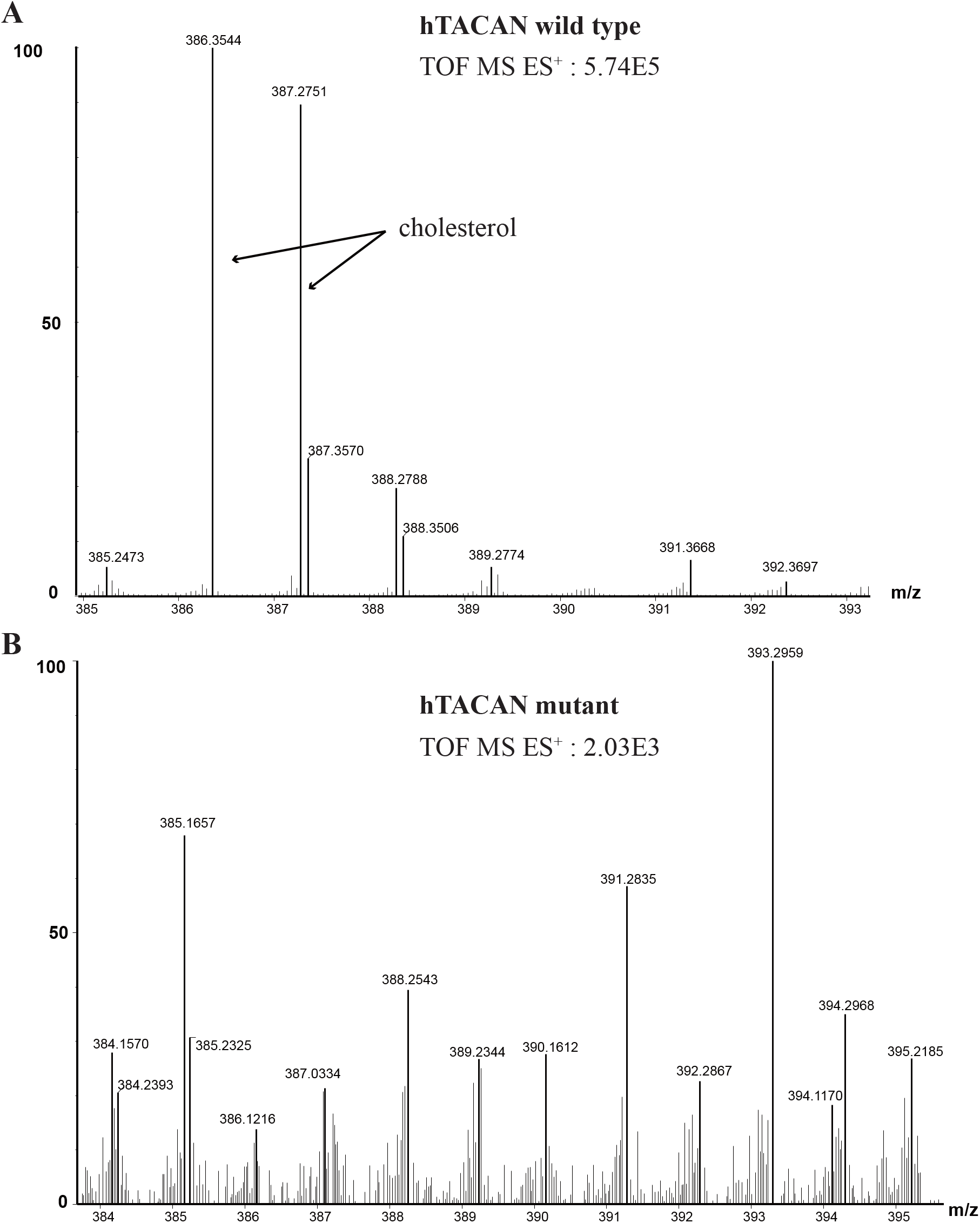
Cholesterol identified in purified hTACAN wild type protein by LC-MS/MS. **A**, Purified hTACAN wild type (0.1 mg) and **B**, mutant F284A-W285A-F288A (0.2 mg) were treated by methanol and chloroform. The extracted cholesterol was dissolved in methanol with formic acid and subjected to LC-MS/MS analysis. The highest m/z value was shown. The cholesterol molecule was identified in the hTACAN wild type but not in mutant.

**Fig. S7.**
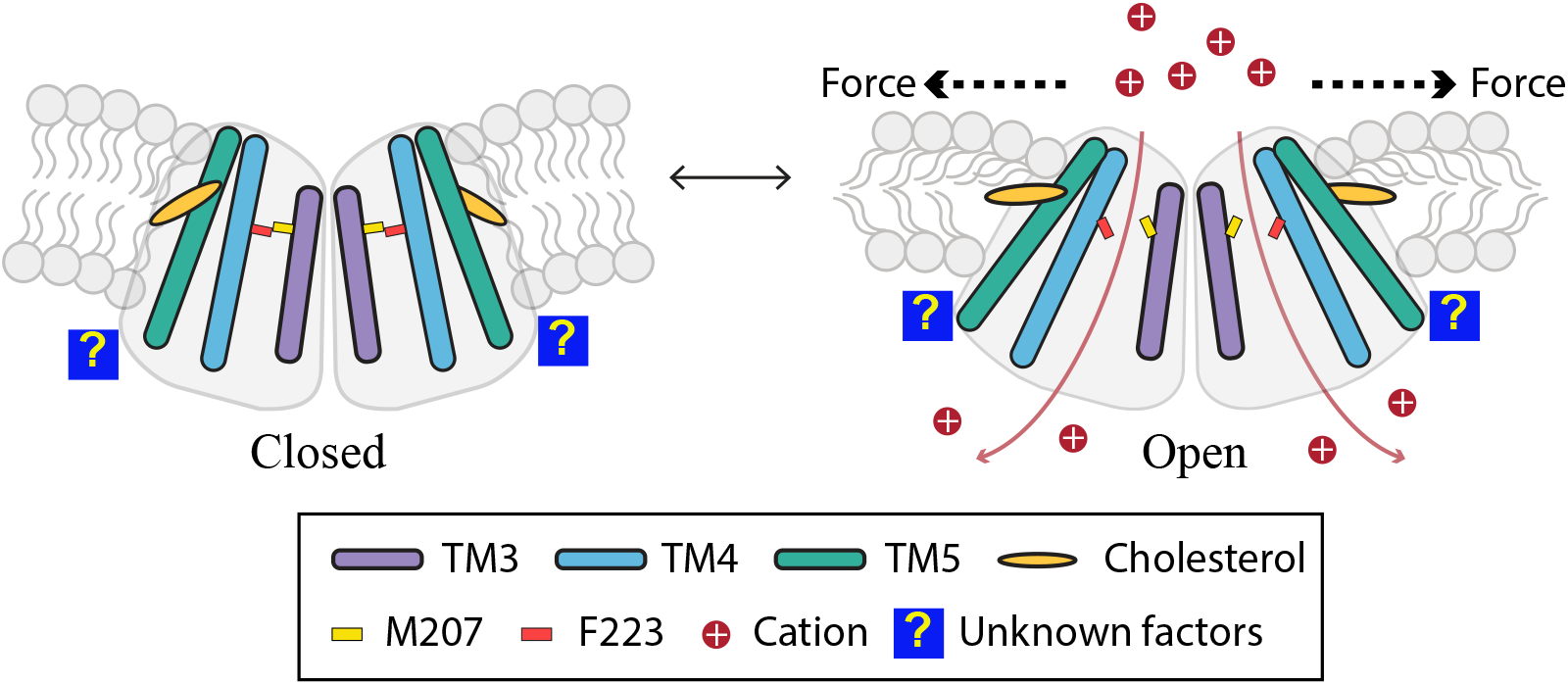
Putative opening mechanism of TACAN under mechanical force. The residue M207 on TM4 and residue F223 on TM5 form a gate and keep the TACAN channel in a closed state in the resting state. When surface tension was sensed by the cholesterols, the movement of TM4 and TM5 resulted in the shifting of M207 and F223 and thus opening the gate. Cations then flow through the channel. Some unknown factors may be required during this process.

